# CRISPR Del/Rei: A simple, flexible, and efficient pipeline for scarless genome editing

**DOI:** 10.1101/2021.01.18.427163

**Authors:** Marah H. Wahbeh, Kyra L. Feuer, Christian Yovo, Eman Rabie, Anh-Thu N. Lam, Sara Abdollahi, Lindsay J. Young, Bailey Rike, Akul Umamageswaran, Dimitrios Avramopoulos

**Author notes:** These authors contributed equally to this work (Marah H. Wahbeh, Kyra L. Feuer). Correspondence should be addressed to: Dimitrios Avramopoulos.

## Abstract

Scarless genome editing of induced pluripotent stem cells (iPSCs) is crucial for the precise modeling of genetic disease. Here we present CRISPR Del/Rei, a two-step deletion-reinsertion strategy with high editing efficiency and simple PCR-based screening that generates isogenic clones in ~2 months. We apply our strategy to edit iPSCs at 3 loci with only rare off target editing.

---

Scarless editing, defined as the introduction of a specific change into the genome without additional mutations, is crucial for modeling disease-associated genotypes. However, doing so with CRISPR-Cas9 is inefficient unless a protospacer or protospacer-adjacent motif (PAM) site is also changed (a “CRISPR-blocking” mutation)^1^. Furthermore, preferred cell types for disease modeling (e.g., iPSCs) usually have low editing efficiency ^2, 3^. While several scarless editing approaches have been described, all have limitations ^4^.

To address these limitations, we used iPSCs to develop CRISPR Deletion and Reinsertion (Del/Rei) (**Fig.1**), an efficient and user-friendly scarless editing strategy. The foundation of CRISPR Del/Rei is the strategic design of three single-guide RNAs (sgRNAs) used in a two-step editing protocol. In Step 1, two sgRNAs recruit Cas9 to create a ~45-110bp deletion that removes a target variant and parts of the protospacers but spares at least one PAM site **(Fig. 1a-b**). The disruption of the protospacers prevents further sgRNA binding and Cas9 cleavage, resulting in high editing efficiency. In Step 2, the deleted sequence is re-inserted (**Fig. 1b-c**) with the desired variant allele(s). The preserved PAM site(s) from Step 1 and the new adjacent sequence is then used as the protospacer for the third sgRNA, which we term “synthetic” (syn-sgRNA). The re-insertion is achieved by homology-directed repair (HDR) using the syn-sgRNA and a single-stranded oligodeoxynucleotide (ssODN) template. Upon reinsertion the syn-sgRNA protospacer is destroyed, once again increasing HDR efficiency.

**Figure 1:**
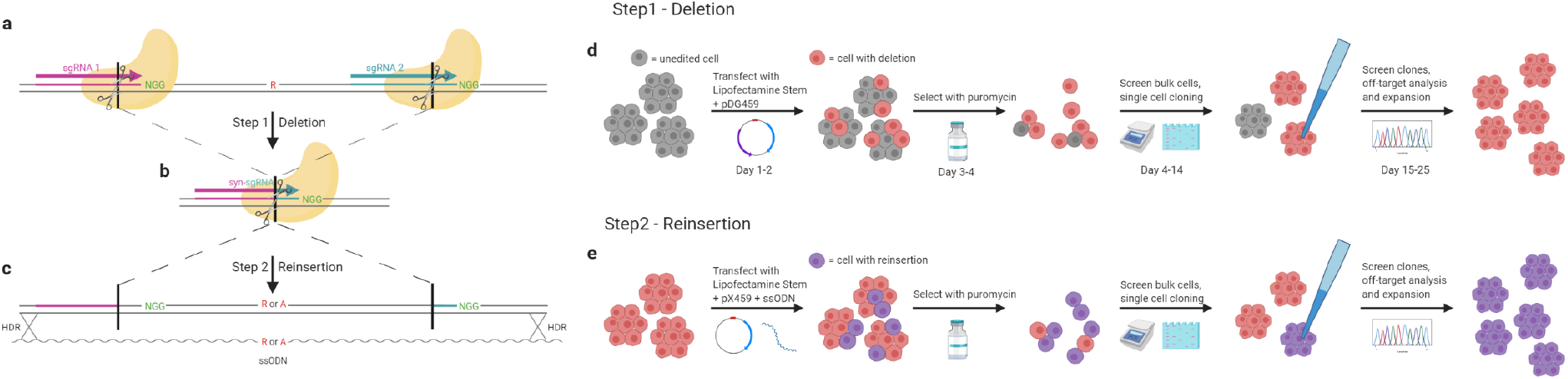
CRISPR Del/Rei strategy: (a-c). experiment design; (d,e). editing process

The CRISPR Del/Rei workflow (**Fig. 2**) is simple and optimized for iPSCs. sgRNAs are cloned in one step ^5^ into the all-in-one vectors pDG459 (Step1) or pX459 (Step 2). iPSCs are plated across multiple wells and transfected with Lipofectamine Stem. Transfected cells undergo positive puromycin selection, which is cheap and reduces the risk of cellular stress, death and contamination compared to cell sorting. After selection, editing efficiency is determined in the “bulk”-cell populations from each transfected well-by simple PCR amplification and electrophoresis. Wells with the highest efficiency are sparse plated for single-cell cloning and again screened for deletions (Step 1) or insertions (Step 2) (**Fig. 2, Supplementary Fig. 1-6)** by PCR-electrophoresis and confirmed by Sanger sequencing. Selected clones are then screened for off-target edits. Different alleles are reinserted in separate transfections or simultaneously through equimolar ssODNs mixes (**Supplementary Fig. 1-4)** The whole process takes ~2 months.

**Figure 2:**
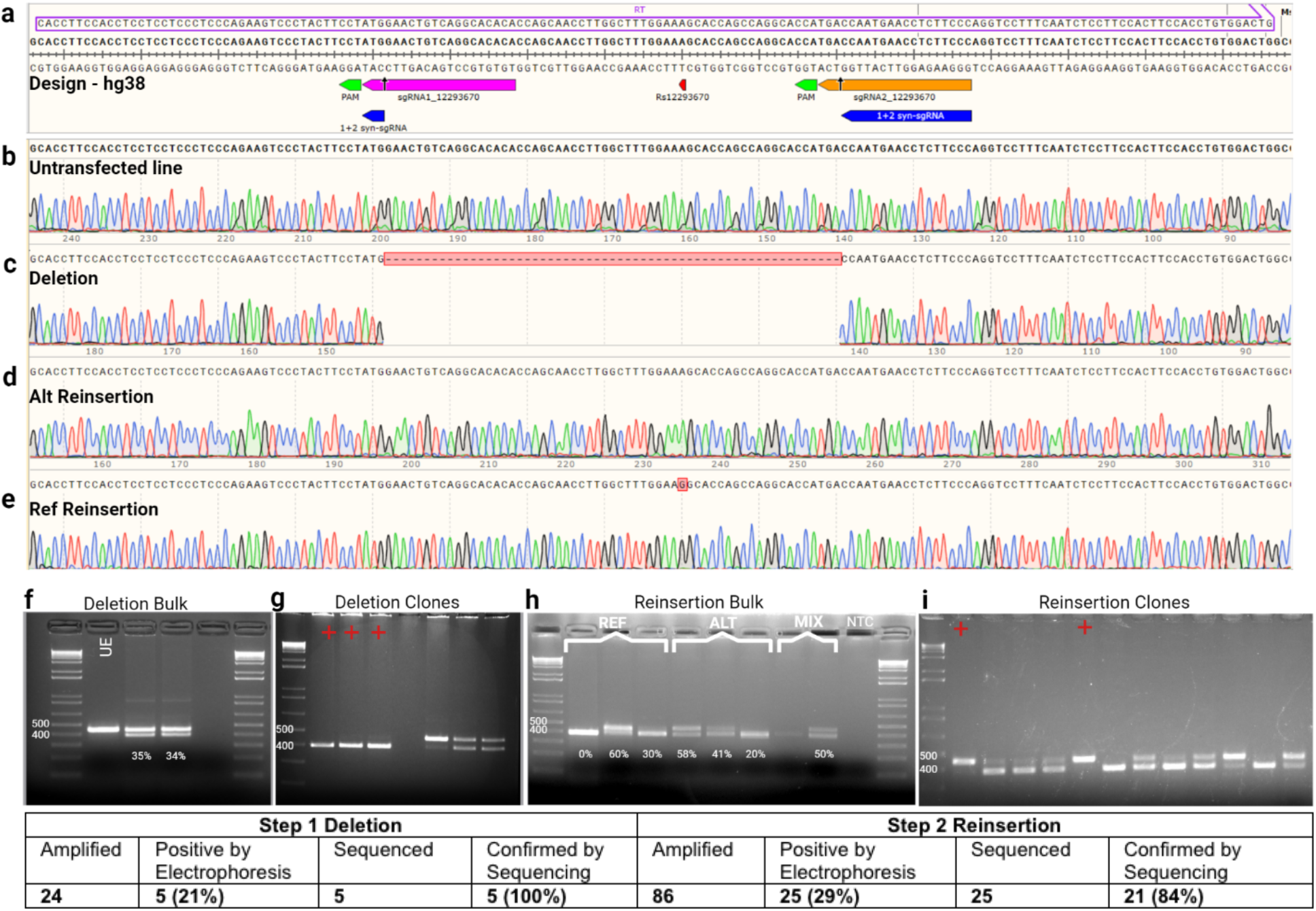
CRISPR Del/Rei example from rs12293670 near *NRGN*: a. Design of gRNAs; b-e sequence confirmations at each step (target base is highlighted); f-I examples of screening by electrophoresis, ladder = 1Kb plus. UE = unedited, NTC = no template control. REF = reference allele template. ALT = alternative allele template. MIX=mixed template.

Using Step 1 we created seven distinct homozygous and one heterozygous deletion (~45-505bp) in 6 schizophrenia-associated coding and noncoding genomic regions across seven different iPSC lines (**Fig 2, Supplementary Figs.1-6, Supplementary Tables 1, 2, and 5**). Average deletion efficiency was 45% (0-100%) with significant variation between technical replicates. Single cell clones found positive by electrophoresis (47%) were confirmed by Sanger sequencing (83%). Prioritized candidate off-target sites were edited at only 5/628 instances across five experiments (**Supplementary Table 3**).

Using Step 2 we performed reinsertions at three loci across 5 different iPSC lines **(Supplementary Tables 1, 2, and 5, Supplementary Figs 1-4).** For each experiment, one off-target-free deletion clone was chosen. Syn-sgRNAs were designed along with 160bp ssODN repair templates for HDR with homology arms flanking the deletion; one ssODN for each variant allele (**Supplementary Table 1**). The two ssODNs were either introduced in separate transfections or simultaneously in an equimolar mix (**Supplementary Figs. 1-4**). Bulk reinsertion efficiency average was 25% (0%-63%, **Fig. 2, Supplementary Figs. 1-4**). After single-cell cloning on average 25% of clones showed re-insertion by electrophoresis (**Fig.** 2, **Supplementary Figs. 1-4**) and sequencing confirmed 60% of those. Mixed ssODNs generated both heterozygous and homozygous clones (**Supplementary Fig 2**). One round of sib-selection ^6^ increased the bulk efficiency and percentage of positive clones up to 100% (**Supplementary Fig. 3 and 4**). Reinsertion clones were screened for off-target editing as Step 1. Off-target editing was detected in only 2 clones from one experiment at one site (**Supplementary Table 3**).

Advantages of CRISPR Del/Rei include high efficiency in iPSCs and identical experimental procedures for the generation of different alleles in isogenic clones. It is flexible in terms of the size and location of the deletion, allowing the targeting of loci that are repetitive or lack PAM sites near the target. Heterozygote edits can also be generated, as well as more complex modifications. When non-coding variants are tested, the deletion can provide a useful first screen for functionality.

Additionally, the transfection and selection protocols are flexible and can be modified to each laboratory’s preferred methods. Strategies to further increase HDR efficiency may be incorporated if desired ^4, 7^. CRISPR Del/Rei was designed and applied in iPSCs, which are difficult to transfect and edit. In more commonly used cancer-derived or immortalized cell lines higher efficiencies are expected.

As with most editing approaches, efficiency varies between transfections, across cell lines, and for different target sites. There is likely increased risk of off-target editing due to the use of 3 sgRNAs, however, our data suggests this is not a significant problem (**Supplementary Table 3**). Additionally, only one deletion clone is required to complete Step 1, therefore off-target effects can be mitigated by selecting an off-target-free clone. Introducing large deletions (>110 bp) is not ideal because the reinsertion would require the use of plasmids as HDR templates, which are less efficient than ssODNs ^8^. Our design requires having 2 PAM sites within 110bp, including one that is external to the cut site, which may be a challenge in some genomic regions but was never a limitation in our experiments.

CRISPR Del/Rei is a novel, effective strategy for quick and efficient scarless editing, that is especially advantageous for generating isogenic cell lines to study disease-associated variants. We believe that in combination with cell differentiation to specific cell types or the generation of organoids, it can provide a significant benefit to future research.

## ACKNOWLEDGEMENTS

This work was supported in part by NIMH grants P50 MH094268, R01 MH113215 and RF1 MH122936 to DA. Figures were created with BioRender.com.

