## Supplementary methods for "CRISPR Del/Rei: A simple, flexible, and efficient pipeline for scarless genome editing"

### Online Methods:

#### Cell lines and cell culture

iPSC lines were obtained from the National Institute of Mental Health (NIMH/RUCDR) stem cell repository (**Supplementary Table 5**, https://www.nimhgenetics.org/). Before cell seeding, tissue culture plates were coated with 5μg/mL laminin (BioLamina #LN521) and incubated at 37℃ for at least two hours. Once seeded, the iPSCs were maintained in a feeder-free culture in StemFlex media with supplement (Gibco #A3349401) at 37℃ with 5% $CO_2$. The media was supplemented with 10 μM ROCK inhibitor (Y-27632 dihydrochloride, Tocris #1254) on the days that thawing, passaging, or transfection took place. Cells were passaged when ~70-80% confluent (~4-5 days) with 1X Phosphate-buffered saline (PBS) and Accutase (MilliporeSigma #A6964). Cells were frozen initially at -80ºC and later in liquid nitrogen in StemFlex media with 10% Dimethylsulfoxide (DMSO). Genotyping of CRISPR targets was done by PCR and Sanger sequencing using primers (25nmole standard desalted, from Integrated DNA Technologies, Inc. [IDT ®] Coralville, IO) with sequences as listed in **Supplementary Table 1** and confirmed by SNP array (see below).

#### Quality Control and authentication of iPSC lines

All iPSC lines were genotyped at the Johns Hopkins University School of Medicine genetics core resource facility on the Illumina Global Screening Array-24 v2.0 to exclude major chromosomal abnormalities and confirm their identity.

#### Design of sgRNAs for deletions (Step 1)

The CRISPOR tool (http://crispor.tefor.net/) was used to identify candidate sgRNAs for each target. sgRNAs were prioritized based on three criteria: location, specificity score, and efficiency score. The location of the sgRNA pair and the resulting deletion they would mediate (**Fig. 2**) was critical for several reasons. First, the deletion was designed such that the target base(s) was near the middle to ensure its removal. Second, the deletion needed to be ~45-110 bp, which is large enough to easily detect by gel electrophoresis, but small enough to accommodate use of an ssODN repair template. Third, at least one of the two PAM sites needed to be external to the deletion to ensure that one was left intact for use by a syn-sgRNA for reinsertion. Fourth, sgRNAs were placed in consideration of any local heterozygous variation. If clones that were homozygous for the target base(s) were desired, sgRNAs were placed to avoid including any heterozygous variants in the deletion. This placement avoided the need to restore such heterozygosity during reinsertion. Finally, in cases where gene knock-out was desired (*DPYSL2* and *NRXN1*) the sgRNAs were placed so the deletion removed known functional domains (i.e. transcription start sites and/or start codons). If heterozygous deletion clones were desired **(Supplementary Fig. 5**) one sgRNA was placed so that a heterozygous SNP was in the seed sequence of the protospacer.

If there were still multiple sgRNA options available after considering the necessary location restrictions, those with high specificity and efficiency scores were preferred. For each variant, multiple sgRNAs were selected and tested as different pairs. The sgRNAs that successfully created deletions and the coordinates of the resulting deletions are listed in **Supplementary Table 1**.

#### Design of syn-sgRNAs and ssODNs for reinsertions (Step 2)

Syn-sgRNAs were designed to match the exact sequence of the corresponding deletion from Step 1. This was determined by Sanger sequencing. They were positioned over the junction of the deletion such that they utilized one of the remaining PAM sites from the sgRNAs (**Fig. 2**). Before ordering the syn-sgRNA, the sequences were queried against the human genome using NCBI BLASTn (https://blast.ncbi.nlm.nih.gov/Blast.cgi) to confirm the absence of identical genomic sites.

The 160bp ssODNs (4 nmol PAGE-purified, IDT) were designed to have homology arms of at least 45bp flanking the deletion from Step 1. Sequences for syn-sgRNAs and ssODN repair templates are listed in **Supplementary Table 1**.

### Cloning

sgRNAs were purchased as single-stranded DNA oligos (25nmoles standard desalted, IDT) with BbsI sticky ends and cloned in pairs into the pDG459 plasmid (Addgene #100901) for the deletion step using a one-step cloning reaction as described by the Thomas lab [25] The same protocol with the minor modification of replacing the second oligo with water during ligation was used to clone the syn-sgRNAs into pX459 (Addgene #62988). A cloning mixture volume of 5μL or 10μL was then transformed into 25μL or 50μL, respectively, of One Shot TOP10 Chemically Competent E.Coli (ThermoFisher #C404003) following the manufacturer's protocol. Transformed bacteria were plated on carbenicillin or ampicillin LB agar plates and incubated at 37℃ overnight. Individual colonies were picked into 3-5mL of LB broth and cultured at 37℃ overnight. Isolation of the plasmid DNA from bacterial clones was conducted using Zymo or Qiagen mini prep and maxi prep kits (Qiagen #27104, Zymo #D4208T)

### Transfection and selection

Transfection and selection parameters differed slightly between experiments due to variability in viability, transfection and editing efficiencies across iPSC lines and genomic loci. The details for each experiment are listed in **Supplementary Table 2**. To summarize, cells were plated 24-48 hours before transfection at densities between 40k-60k cells/well on 24-well plates coated with laminin. The cells were transfected with 500ng of pDG459 (Step 1) or pX459 and ssODN templates at a 1:20 molar ratio using 2μL of Lipofectamine Stem (ThermoFisher #STEM00003) per well. ssODN templates for different alleles were added together in the same transfection for some experiments allowing for generation of homozygous and heterozygous clones from one transfection while others were added in parallel in separate transfection wells. We followed the published Lipofectamine Stem protocol (https://assets.thermofisher.com/TFS-Assets/BID/manuals/transfection-psc-lipofectamine-stem-stemflex-protocol.pdf), sometimes with alterations. Specifically, in some cases the cells were incubated in the DNA/Lipofectamine/Opti-MEM mixture for 5-7 hours instead of the recommended 4 hours because this increased the editing efficiency. Additionally, after the incubation period the DNA/Lipofectamine/Opti-MEM mixture for some cell lines was exchanged with StemFlex instead of being supplemented with StemFlex because this increased cell viability. Puromycin was added ~24-48 hours post transfection at optimal concentration for each cell line. The cells were left to recover for 3-7 days before passaging and screening for bulk editing efficiency.

### Screening and expansion of edited cells

To screen for desired edits, genomic DNA from bulk cells of each transfected well was extracted using Quick Extract buffer (Lucigen #QE0905T). following the protocol published by the manufacturer (https://www.lucigen.com/docs/manuals/MA150E-QuickExtract-DNA-Solution.pdf). Briefly, ~5-7 days after transfection media was collected from the transfected wells and spun down. Small pellets from floating/ dead cells were detected and the volume of Quick Extract solution volume added (10-50μL) was determined by how large the pellet was. This allowed for a quick screen of the bulk cells. If amplification using Quick Extract-derived DNA was not successful, then cells were expanded so a larger cell pellet could be obtained, and the Gentra Puregene Cell Kit was used for extraction instead (Qiagen #158388). Amplicons ~200-850bp flanking the target site (**Supplementary Table 1**) were amplified using 5μL (Quick Extract) or 1μL (Gentra Puregene) of extracted DNA and the AccuPrime Taq DNA Polymerase System (ThermoFisher #12339016). A volume of 10-15μL of each PCR product was combined with 10X Bromophenol Blue loading dye and loaded into 1-2% agarose gels and run at 80-120V for 30-60 minutes. The gel percentage, voltage and running time depended on the size of the amplicons and the expected deletion. Samples that appeared positive for editing by gel were quantified using image J and sent for Sanger sequencing to confirm the exact edited sequence.

Transfection wells with the highest editing efficiency underwent single-cell cloning, which consisted of sparsely plating 300-500 cells per well in 6-well plates (2mL of media per well) and allowing colonies to develop over 7-10 days. Half of each colony was manually picked into a PCR tube in ~10μL of media and the other half into one well of a 24- or 96-well culture plate. The half clones in PCR tubes were mixed with 10μL of Quick Extract buffer for DNA extraction, followed by PCR, gel electrophoresis, and Sanger sequencing to confirm genotype. Positive clones were then expanded for off-target analysis.

If reinsertion efficiency was lower than desired, we incorporated one or a combination of the following strategies: cold shocking the cells by placing them in a 32℃ with 5% $CO_2$ incubator for 48 hours after transfection and sib-selection. Sib selection was performed to generate populations of cells enriched for the edit. This consisted of passaging bulk transfected cells into 96-well plates at 30 cells/well. Once the wells reached confluency, the media was harvested and spun down to obtain cell pellets. Pellets were mixed with 10-30μL of Quick Extract, and screening proceeded as described above. Sib-selection wells that showed enrichment for a reinsertion edit were then sparse plated for single-cell cloning, from which individual colonies were picked and screened.

**Off-target analysis**

All predicted off-target sites for each sgRNA and syn-sgRNA were obtained from the CRISPOR tool (http://crispor.tefor.net/). To narrow down the number of sites to those more likely, we prioritized sites with 3 or less mismatches with our sgRNAs and among these sites most likely to be in functional sequence either because they were in coding sequence or in DNase1 hypersensitivity sites (DHS) obtained from ENCODE (https://www.encodeproject.org/, file: hg38_wgEncodeRegDnaseClustered). If a small number of candidate off-target sites was identified, we included additional sites with up to 4 mismatches and/or those within 100-500 bp of a DHS. Individual clones with confirmed edits For Step 1 and Step 2 were tested for off-target edits by Sanger sequencing of prioritized off-target sites and compared to the un-transfected line. All off-target sites and primers used are summarized in **Supplementary Tables 3 and 4**.
