## Supplementary Tables and figures for "CRISPR Del/Rei: A simple, flexible, and efficient pipeline for scarless genome editing"

**Supplementary Figure 1 | CRISPR Del/Rei editing of rs4129585 (*TSNARE1*) in RU17 iPSC line**.
**(a)** sgRNA, syn-sgRNA, and ssODN repair template design. **(b-f)** Sanger sequencing chromatograms of the cell line before and after each step of editing. **(g)** Screen of the bulk deletion transfection cells. Each lane represents a technical replicate from a separate transfection well. Editing efficiency quantified by Image J shown under each lane. Non-control lanes without an indicated percentage were from a different experiment and can be ignored. The lane marked with a green arrow was used for single cell cloning. (**h)** Screen of deletion clones. Each lane represents one clone. Lanes with a red plus were confirmed positive by Sanger sequencing. **(i)** Screen of the bulk reinsertion transfection cells. The reference and alternate alleles were reinserted in parallel in separate wells (REF and ALT), and simultaneously in the same wells (MIX). **(j)** Representative screen of the mix reinsertion clones. (**k)** Summary of total clones screened for each experiment. The positive by electrophoresis percentage refers to the number of clones that looked positive by gel out of the total clones amplified. The confirmed by sequencing percentage refers to the number of clones that were confirmed positive by Sanger sequencing out of the total number of clones sent for sequencing. All ladders are 1kb plus. DEL = deletion positive control (402 bp in deletion bulk and clone gels, using F2 and R2 primers and 362 bp in reinsertion bulk and clone gels using R1 and R2 (**Sup. Table 1)** , UE = unedited positive control 447 bp in deletion bulk and clone gels, using F2 and R2 primers and 407 bp in reinsertion bulk and clone gels using R1 and R2(**Sup. Table 1**), NTC = no template negative control.

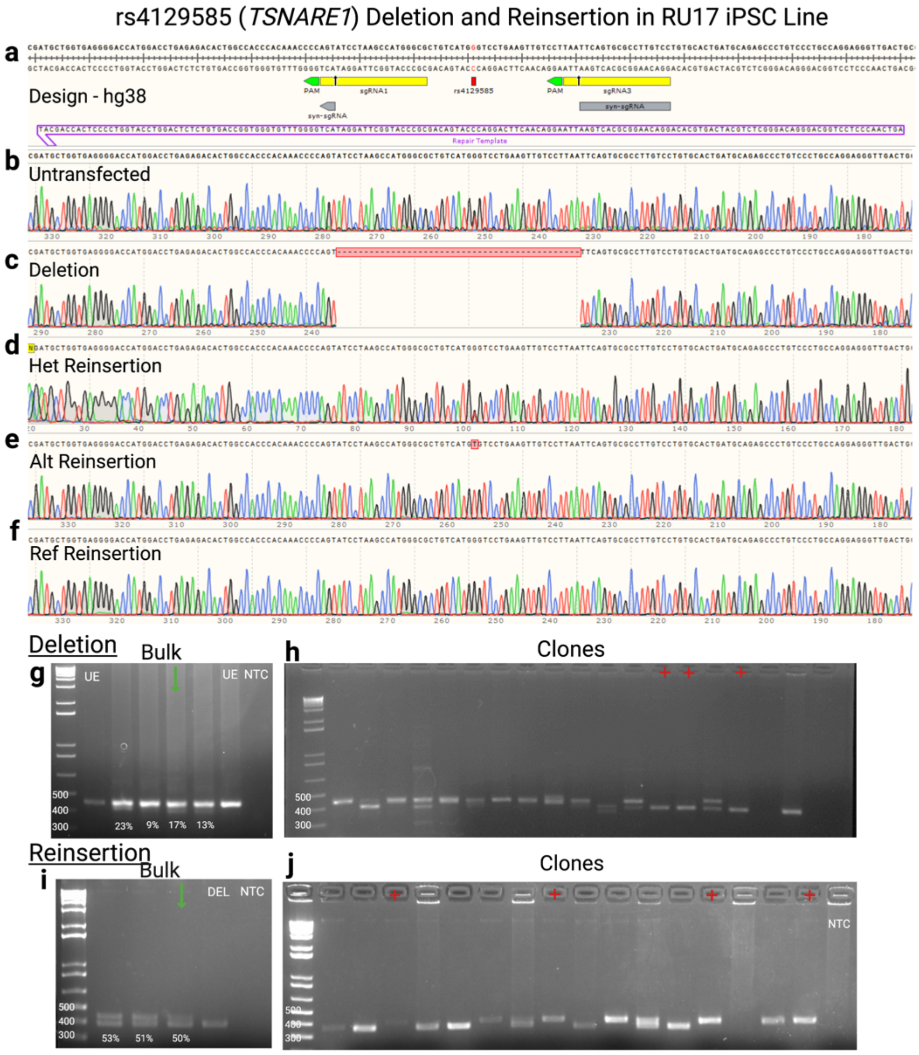


**k**

| Step 1 Deletion | | | | Step 2 Reinsertion | | | |
|---|---|---|---|---|---|---|---|
| Amplified | Positive by Electrophoresis | Sequenced | Confirmed by Sequencing | Amplified | Positive by Electrophoresis | Sequenced | Confirmed by Sequencing |
| **17** | **5 (29%)** | **4** | **2 (50%)** | **131** | **24 (18%)** | **12** | **12 (100%)** |

**Supplementary Figure 2 | CRISPR Del/Rei editing of rs4129585 (*TSNARE1*) in the RU04 iPSC line.**
**(a)** sgRNA, syn-sgRNA, and ssODN repair template design. **(b-f)** Sanger sequencing chromatograms of the cell line before and after each step of editing. **(g)** Screen of the bulk deletion transfection cells. Each lane represents a technical replicate from a separate transfection well. . Editing efficiency quantified by Image J shown under each lane. Non-control lanes without an indicated percentage were from a different experiment and can be ignored. The lane marked with a green arrow was used for single cell cloning. **(h)** Screen of deletion clones. Each lane represents one clone. Lanes with a red plus were confirmed positive by Sanger sequencing. **(i)** Screen of the bulk reinsertion transfection cells. The reference and alternate alleles were reinserted in parallel in separate wells (REF and ALT), and simultaneously in the same wells (MIX). **(j)** Representative screen of the mix reinsertion clones. (**k**) Summary of total clones screened for each experiment. The positive by electrophoresis percentage refers to the number of clones that looked positive by gel out of the total clones amplified. The confirmed by sequencing percentage refers to the number of clones that were confirmed positive by Sanger sequencing out of the total number of clones sent for sequencing.
All ladders are 1kb plus. Primers used: Forward 2 and Reverse 2, see Supplement table 1. DEL = deletion positive control 402 bp, UE = unedited positive control 447 bp, NTC = no template negative control.

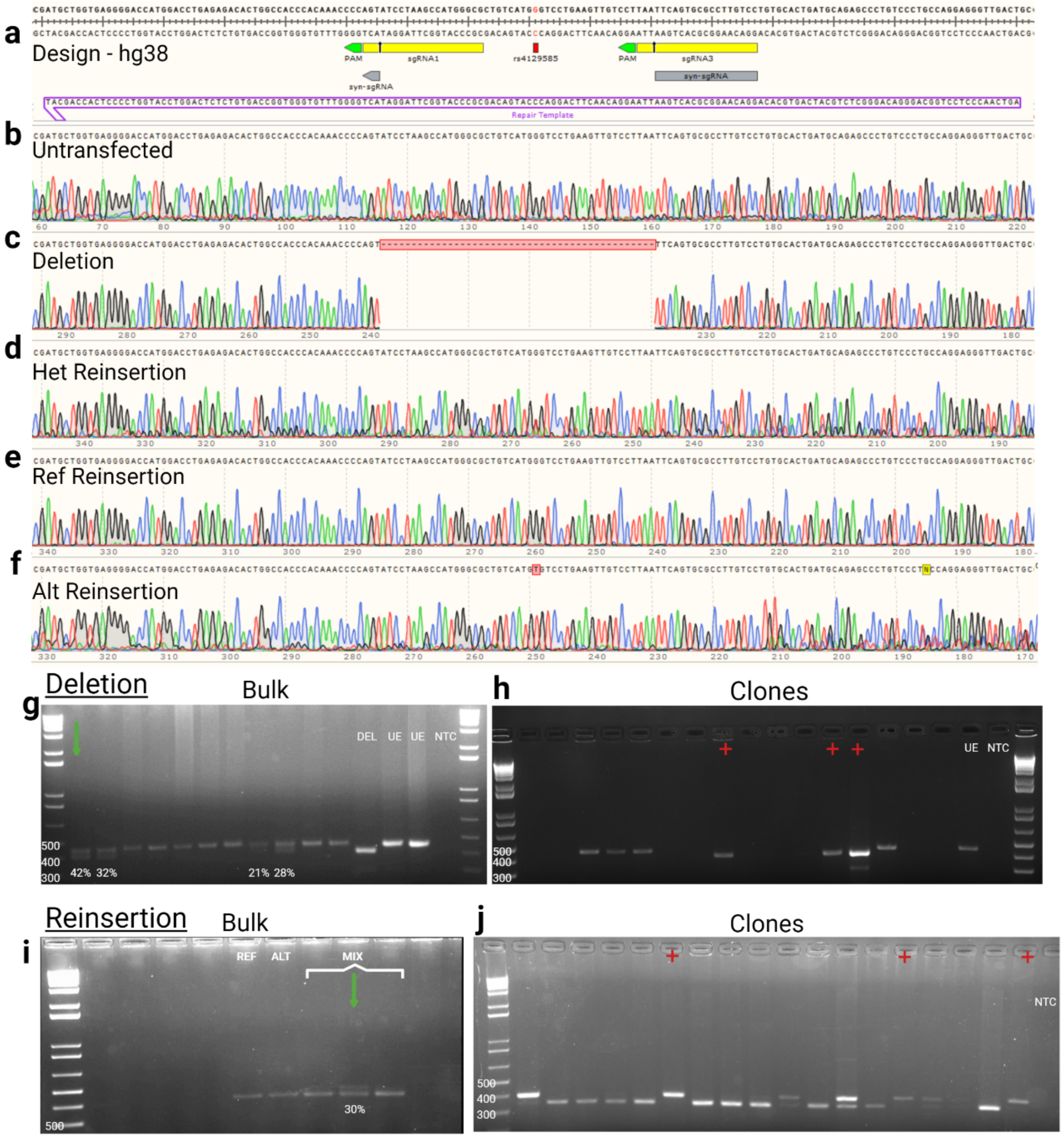


**k**

| Step 1 Deletion | | | | Step 2 Reinsertion | | | |
|---|---|---|---|---|---|---|---|
| Amplified | Positive by Electrophoresis | Sequenced | Confirmed by Sequencing | Amplified | Positive by Electrophoresis | Sequenced | Confirmed by Sequencing |
| **12** | **5 (42%)** | **2** | **2 (100%)** | **118** | **20 (17%)** | **20** | **20 (100%)** |

**Supplementary Figure 3 | CRISPR Del/Rei of rs4766429 (*ATP2A2*) in the RU07 iPSC Line.**
**(a)** sgRNA, syn-sgRNA, and ssODN repair template design. **(b-e)** Sanger sequencing chromatograms of the cell line before and after each step of editing. **(f)** screen of the bulk cells from each deletion transfection well. Each lane represents a technical replicate from a separate transfection well unless otherwise marked. Editing efficiency quantified by Image J shown under each lane. The lane marked with a green arrow was expanded for single cell cloning. **(g)** Example deletion clone screening gel. Each lane represents one clone. Lanes marked with a red cross were positive clones confirmed by Sanger sequencing. The reference (ref) and alternate (alt) alleles were reinserted in sequential transfections. **(h)** screen of the bulk cells from the alt reinsertion transfection. The lane marked with the green arrow was expanded for the single cell cloning and for sib selection. **(i)** example screen of the alternate allele clones derived from single-cell cloning. **(j)** sib-selection bulk screen. Each lane represents a population of cells grown in one well of a 96-well plate. Cells from the lane marked with a green arrow were expanded for single cell cloning. **(k)** example sib selection clone screening gel. **(l)** screen of the bulk cells from the ref reinsertion transfection. Cells from the lane marked with the green arrow were expanded for sib selection. **(m)** sib selection bulk screens. **(n)** sib selection clones. **(o)** Summary of total clones screened for each experiment. The positive by electrophoresis percentage refers to the number of clones that looked positive by gel out of the total clones amplified. The confirmed by sequencing percentage refers to the number of clones that were confirmed positive by Sanger sequencing out of the total number of clones sent for sequencing.
All ladders are 1Kb plus. UE = unedited/reinsertion positive control 578bp, NTC = no template negative control, DEL = deletion positive control 511bp.

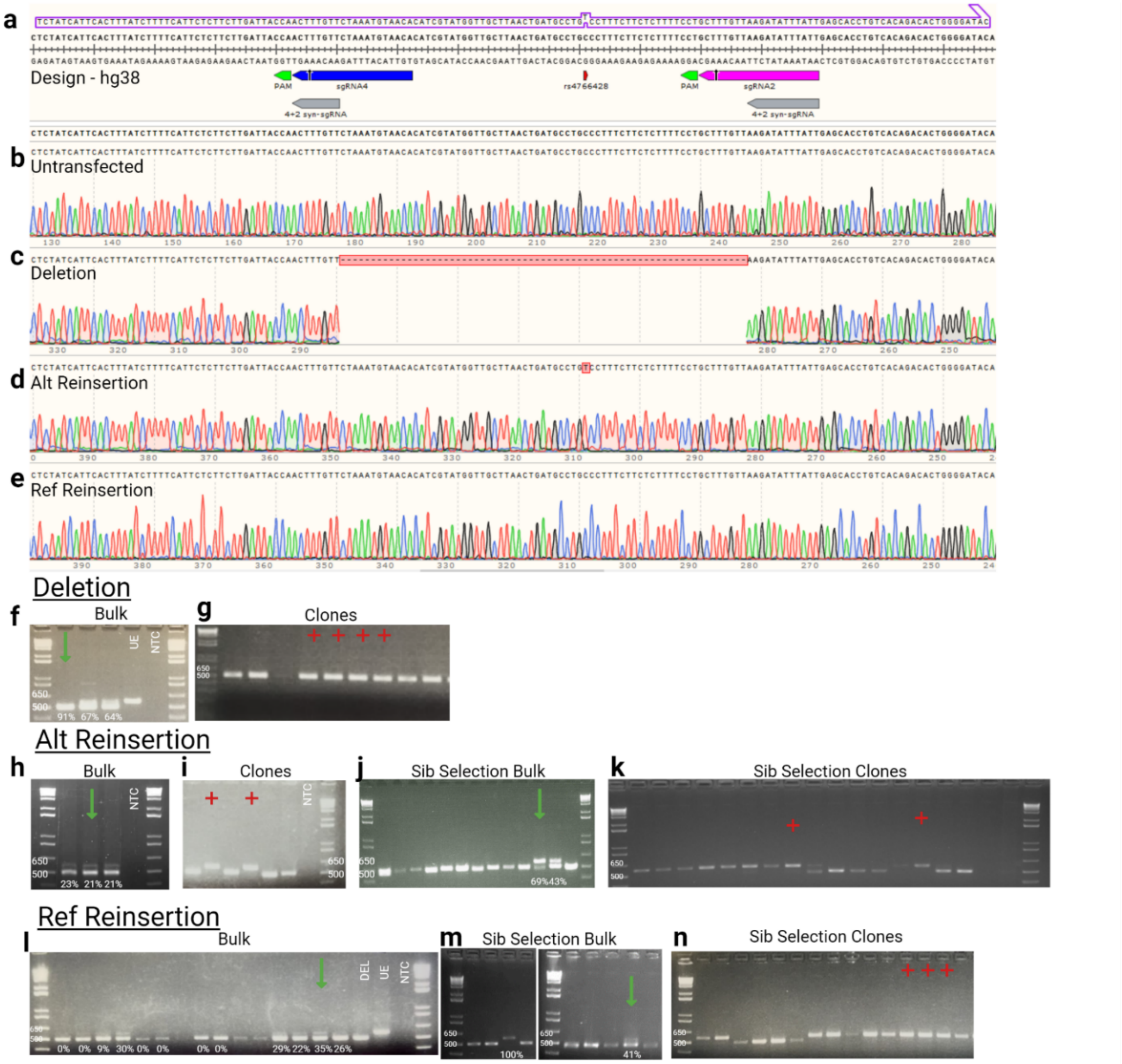


**o**

| Step 1 Deletion | | | | Step 2 Reinsertion | | | |
|---|---|---|---|---|---|---|---|
| Amplified | Positive by Electrophoresis | Sequenced | Confirmed by Sequencing | Amplified | Positive by Electrophoresis | Sequenced | Confirmed by Sequencing |
| **24** | **24 (100%)** | **5** | **5 (100%)** | **146** | **56 (38%)** | **25** | **19 (76%)** |

**Supplementary Figure 4 | CRISPR Del/Rei of rs4766429 (*ATP2A2*) in the RU02 iPSC Line.**
(**a**) sgRNA, syn-sgRNA, and ssODN repair template design. **(b-d)** Sanger sequencing chromatograms of the cell line before and after each step of editing. **(e)** screen of the bulk cells from each deletion transfection well. Each lane represents a technical replicate from a separate transfection well unless otherwise marked. Editing efficiency quantified by Image J shown under each lane. The lane marked with a green arrow was expanded for single cell cloning. **(f)** Example clone screening gel. Each lane represents one clone. Lanes marked with a red cross were positive clones confirmed by Sanger sequencing. **(g)** screen of the bulk cells from reinsertion transfections. The reference and alternate alleles were reinserted in parallel in separate wells (REF and ALT), and simultaneously in two wells (MIX). The ALT lane marked with the green arrow was expanded for the single cell cloning. **(h)** example screen of the alternate allele clones. **(i)** Summary of total clones screened for each experiment. The positive by electrophoresis percentage refers to the number of clones that looked positive by gel out of the total clones amplified. The confirmed by sequencing percentage refers to the number of clones that were confirmed positive by Sanger sequencing out of the total number of clones sent for sequencing.
All ladders are 1Kb plus. UE = unedited/reinsertion positive control 578bp, NTC = no template negative control, DEL = deletion positive control 511bp.

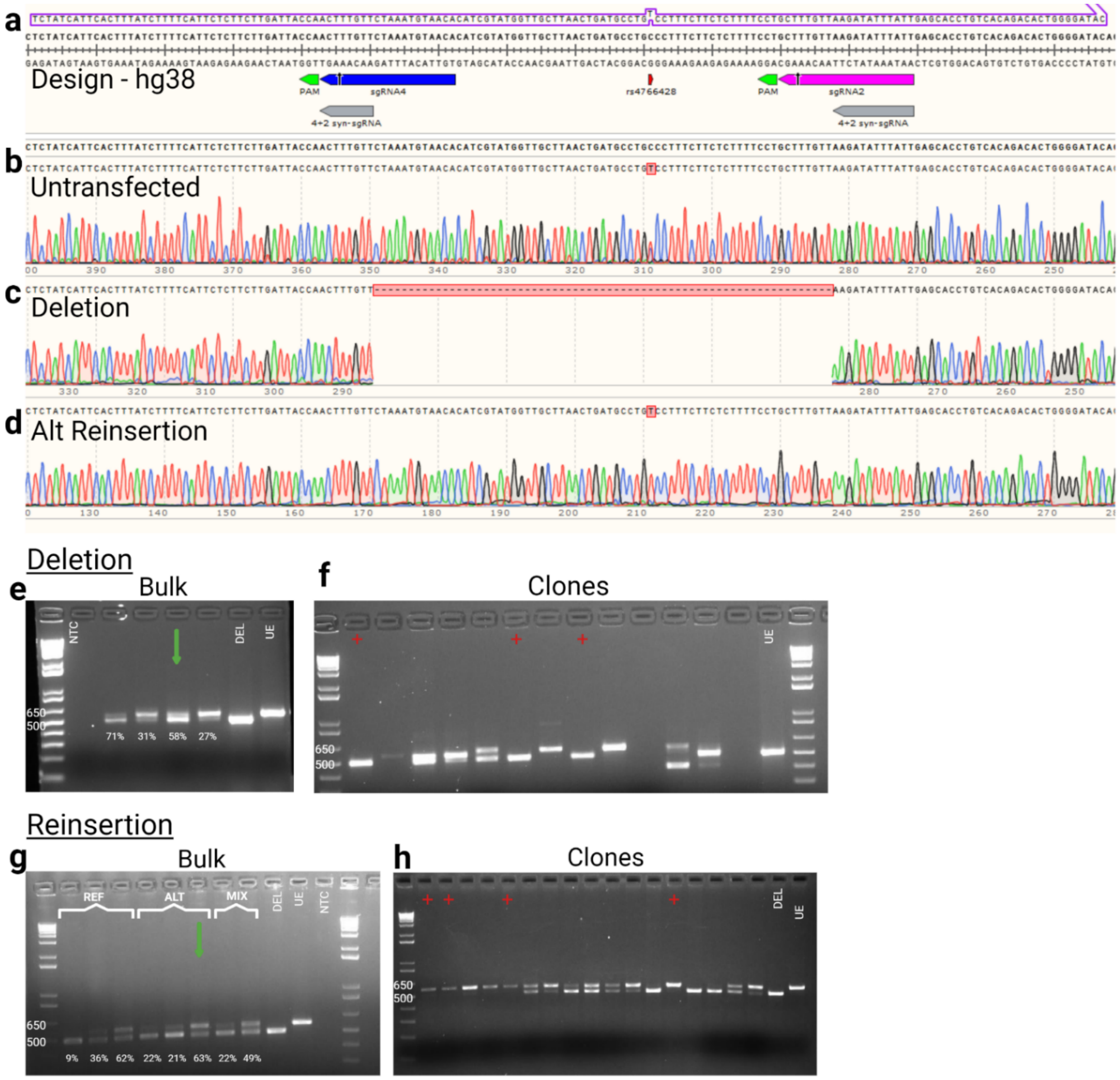


i

| Step 1 Deletion | | | | Step 2 Reinsertion | | | |
|---|---|---|---|---|---|---|---|
| Amplified | Positive by Electrophoresis | Sequenced | Confirmed by Sequencing | Amplified | Positive by Electrophoresis | Sequenced | Confirmed by Sequencing |
| **15** | **6 (40%)** | **6** | **4 (67%)** | **79** | **12 (15%)** | **12** | **11 (92%)** |

**Supplementary Figure 5 | Generating heterozygotes deletions of rs4766428 (*ATP2A2*) in the RU05 iPSC line using step 1 of CRISPR Del/Rei.**
 (**a**) CRISPR Design (top panel). The cell line used for this experiment (bottom panel) was heterozygous for a common SNP (rs3026445) located in the seed sequence of sgRNA 4. **(b)** Screen of the bulk transfected deletion cells. Each lane represents a technical replicate from a separate transfection well unless otherwise marked. Editing efficiency quantified by Image J shown under each lane. The lane marked with a green arrow was expanded for single cell cloning. **(c)** Screen of single-cell derived clones. Of the 41 clones that successfully amplified, 29 (71%) appeared to be heterozygous by electrophoresis and our inability to separate pure deletion clones by repeated sparse plating and clone selection experiments. All ladders are 1Kb plus. UE = unedited positive control (578bp), NTC = no template negative control, Expected deletion size = 511bp.

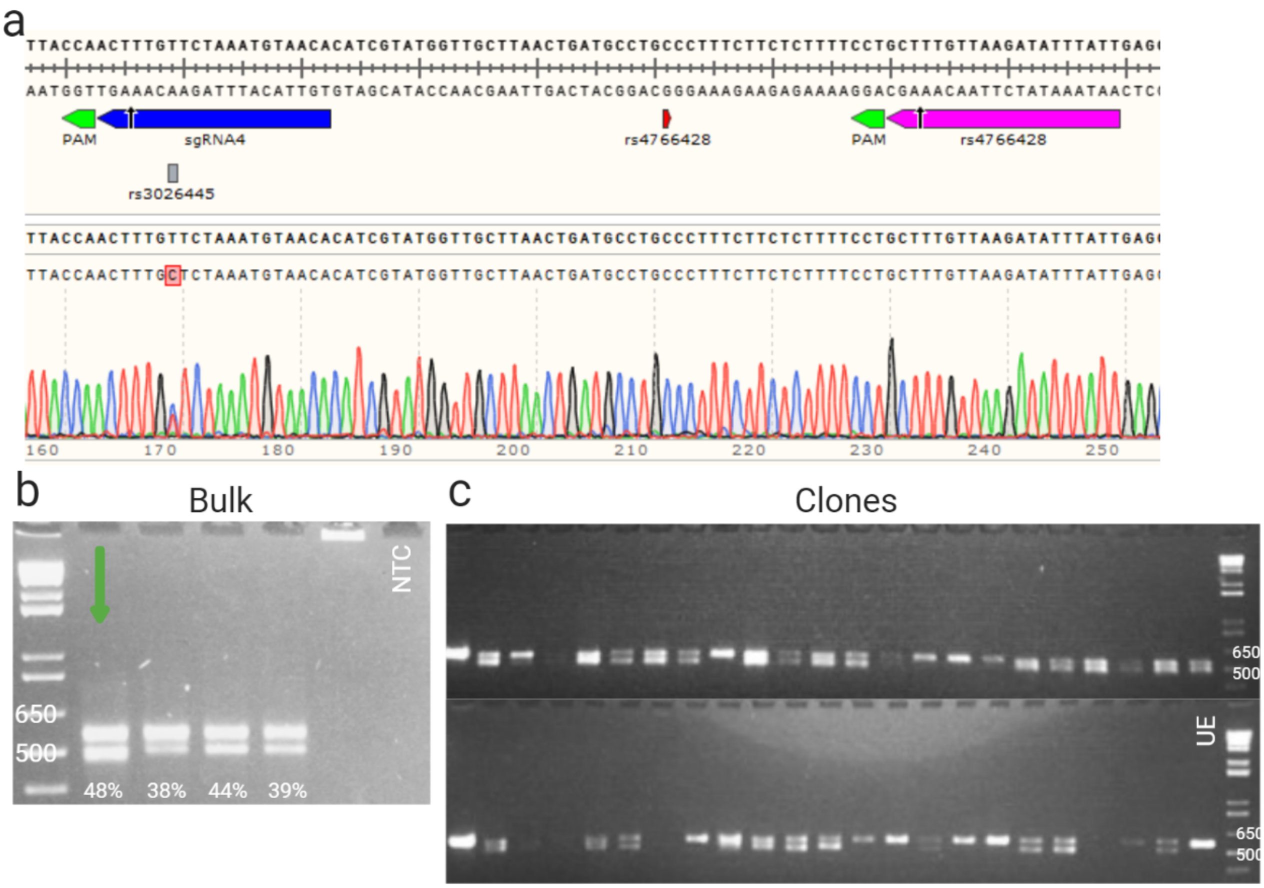

**Supplementary Figure 6 | CRISPR design, chromatograms, gels, and clone information for deletions targeting rs4766428 (*ATP2A2*), *NRXN1, DPYSL2*, and *PKNOX1*.** (**a,e,i,m)** CRISPR design based on hg38, and chromatograms of the untransfected lines and one example deletion clone for each experiment. (**b,f,j,n)** Screens of the bulk transfected cells from each experiment. Each lane represents a technical replicate from a separate transfection well unless otherwise marked. Lanes marked with a green arrow were expanded for single cell cloning **(c,g,k,o)** Example clone screening gels from each experiment. Each lane represents one clone. Lanes marked with a red cross were positive clones confirmed by Sanger sequencing. (**d,h,l,p)** Summary of total clones screened for each experiment. The positive by electrophoresis percentage refers to the number of clones that looked positive by gel out of the total clones amplified. The confirmed by sequencing percentage refers to the number of clones that were confirmed positive by Sanger sequencing out of the total number of clones sent for sequencing. Editing efficiency quantified by Image J shown under each lane.
All ladders are 1Kb+. UE = unedited positive control, NTC = no template negative control.

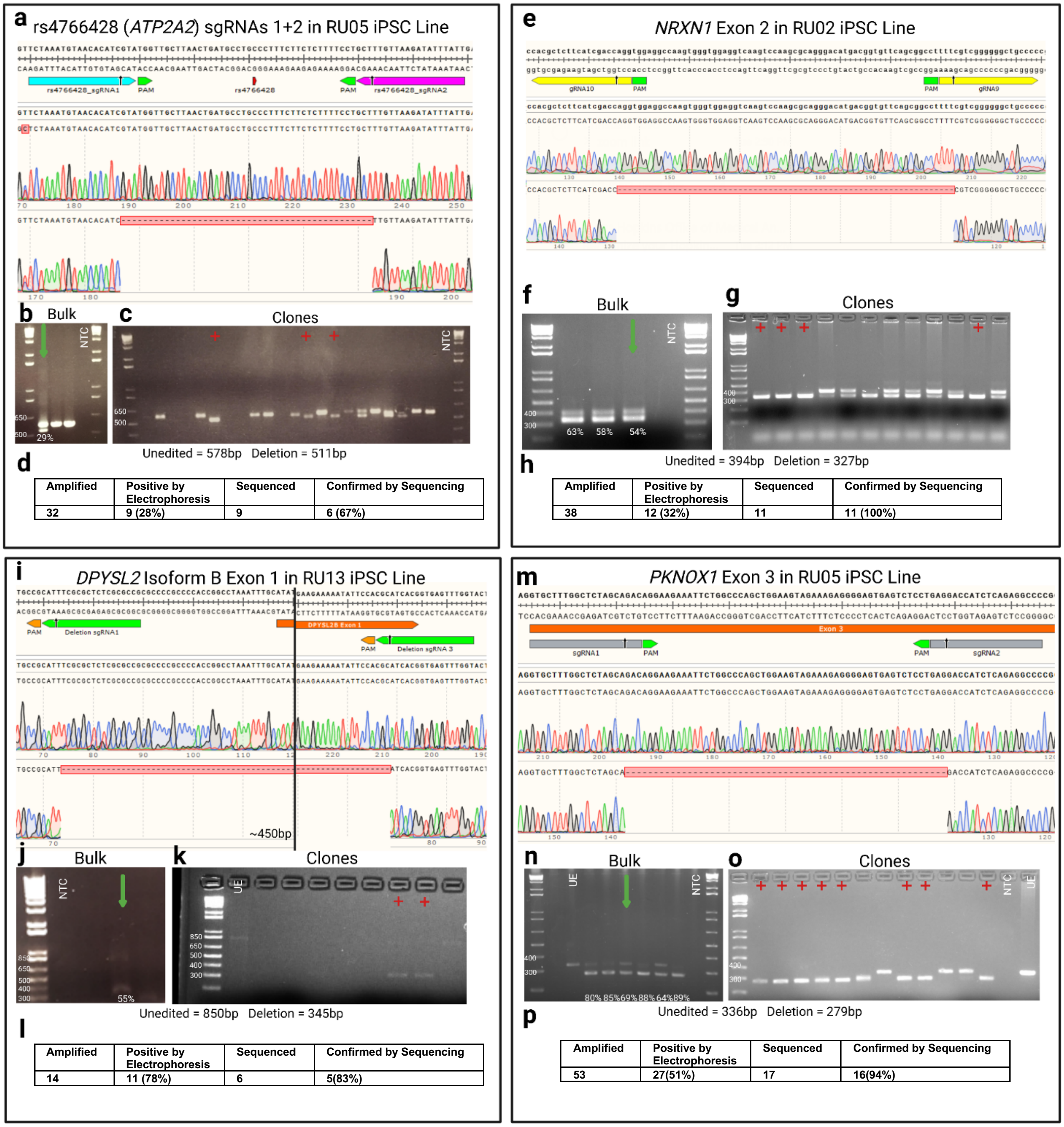


d

| Amplified | Positive by Electrophoresis | Sequenced | Confirmed by Sequencing |
|---|---|---|---|
| 32 | 9 (28%) | 9 | 6 (67%) |

h

| Amplified | Positive by Electrophoresis | Sequenced | Confirmed by Sequencing |
|---|---|---|---|
| 38 | 12 (32%) | 11 | 11 (100%) |

l

| Amplified | Positive by Electrophoresis | Sequenced | Confirmed by Sequencing |
|---|---|---|---|
| 14 | 11 (78%) | 6 | 5(83%) |

p

| Amplified | Positive by Electrophoresis | Sequenced | Confirmed by Sequencing |
|---|---|---|---|
| 53 | 27(51%) | 17 | 16(94%) |

### Supplementary Table 1 | Target sites CRISPR details for each experiment

| Locus | Target(s) | Genotyping and Sequencing Primers | Deletion Coordinates (GRCh38/hg38) | Deletion sgRNAs and PAMs | Reinsertion syn-sgRNAs and PAM |
|---|---|---|---|---|---|
| ***TSNARE1*** | **rs4129585** | **Forward 1: 5'-GGGGGATGATGCTTGGGAAA-3'**<br>**Reverse 1: 5'-GTGGCAGATCTGAGGTCCAG-3'**<br>**Forward 2: 5'-TAGACTGGGGGTGTTGGTGG-3'**<br>**Reverse 2: 5'- AGACCTTCCCAGGCTTCACAG-3'** | **chr8:142231553-142231597** | **sgRNA1: 5'-CGCCCATGGCTTAGGATACT GGG-3' (-)**<br>**sgRNA3: 5'-GGACAAGGCGCACTGAATTA AGG-3' (+)** | **5'-GGACAAGGCGCACTGAAACT GGG-3' (-)** |
| ***NRGN*** | **rs12293670** | **Forward: 5'-TAGCACAGGGCCTGACATAAT-3'**<br>**Reverse: 5'-GAATGGAAACCTGGAATAGCA-3'** | **chr11:124742998-124743056** | **sgRNA1: 5'-TGTGTGCCTGACAGTTCCAT AGG-3'**<br>**sgRNA2: 5'-GGGAAGAGGTTCATTGGTCA TGG'-3** | **5'-ATGCCAATGAACCTCTTCCC cgg-3' (-)** |
| ***ATP2A2*** | **rs4766428** | **Forward: 5'-CTATGCTTGTGACCTGTAGGGA-3'**<br>**Reverse: 5'-AACGACGGTGGAAGGTAGAGA-3'** | **sgRNAs 4 and 2- chr12:110285400-110285466** | **sgRNA2: 5'-AATAAATATCTTAACAAAGC AGG-3' (-)**<br>**sgRNA4: 5'-TGTTACATTTAGAACAAAGT TGG-3' (-)** | **5'-AATAAATATCTTAACAAAGT TGG-3' (-)** |
| | | | **sgRNAs 1 and 2- chr12:110285415-110285461** | **sgRNA1: 5'-TCTAAATGTAACACATCGTA TGG-3' (+)**<br>**sgRNA2: 5'-AATAAATATCTTAACAAAGC AGG-3' (-)** | **N/A** |
| ***NRXN1*** | **Exon 2** (ENSE00002171876) | **Forward: 5'- CGACTTCCTGGAGCTGATTC - 3'**<br>**Reverse: 5'-CTCATCGTCCAGCTTCACCT - 3'** | **sgRNAs 10 and 9- chr2:51027813- 51027879** | **sgRNA10: 5'-CACGCTCTTCATCGACCAGG TGG-3' (+)**<br>**sgRNA9: 5'- GGGGCAGCCCCCCGACGAAA AGG-3'(+)** | **N/A** |
| ***DPYSL2*** | **Exon 1**<br>(ENSE00002094501) | **Forward: 5'-ATTTGCATATCCCAGGATCG-3'**<br>**Reverse: 5'-GCTACCTTCGTGCACTCG-3'** | **chr8:26577780-26578287** | **sgRNA1: 5'-CGGCGCGAGAGCGCGAAATG CGG-3' (-)**<br>**sgRNA3: 5'-ACCAAACTCACCGTGATGCG TGG-3' (-)** | **N/A** |
| ***PKNOX2*** | **Exon 3** (ENSE00002153508) | **Forward: 5'-CTGGAGAGGGGAGAACACAG-3'**<br>**Reverse: 5'-TGATTGGCTTTCAAGGTCGT-3'** | **chr11:125201618+125201953** | **sgRNA1: 5'-GTGCTTTGGCTCTAGCAGAC AGG-3'**<br>**sgRNA2: 5'-GGGCCTCTGAGATGGTCCTC AFF-3'** | **N/A** |

**ssODNs:**

***TSNARE1:***

Alternate: 5'- AGTCAACCCTCCTGGCAGGGACAGGGCTCTGCATCAGTGCACAGGACAAGGCGCACTGAATTAAGGACAACTTCAGGAC**A**CATGACAGCGCCCATGGCTTAGGATACTGGGGTTTGTGGGTGGCCAGTGTCTCTCAGGTCCATGGTCCCCTCACCAGCAT -3'

' Reference: 5'- AGTCAACCCTCCTGGCAGGGACAGGGCTCTGCATCAGTGCACAGGACAAGGCGCACTGAATTAAGGACAACTTCAGGAC**C**CATGACAGCGCCCATGGCTTAGGATACTGGGGTTTGTGGGTGGCCAGTGTCTCTCAGGTCCATGGTCCCCTCACCAGCAT -3'

***ATP2A2:***

Alternate*:* 5'- TCTATCATTCACTTTATCTTTTCATTCTCTTCTTGATTACCAACTTTGTTCTAAATGTAACACATCGTATGGTTGCTTAACTGATGCCTG**T**CCTTTCTTCTCTTTTCCTGCTTTGTTAAGATATTTATTGAGCACCTGTCACAGACACTGGGGATAC - 3'

Reference: 5'- TCTATCATTCACTTTATCTTTTCATTCTCTTCTTGATTACCAACTTTGTTCTAAATGTAACACATCGTATGGTTGCTTAACTGATGCCTG**C**CCTTTCTTCTCTTTTCCTGCTTTGTTAAGATATTTATTGAGCACCTGTCACAGACACTGGGGATAC- 3'

***NRGN:***

Alternate: 5'- CACAGGTGGAAGTGGAAGGAGATTGAAAGGACCTGGGAAGAGGTTCATTGGTCATGGTGCCTGGCTGGTGCCTTCCAAAGCCAAGGTTGCTGGTGTGTGCCTGACAGTTCCATAGGAAGTAGGGACTTCTGGGAGGGAGGAGGAGGTGGAAGGTGCCCTG – 3'

Reference: 5'- CACAGGTGGAAGTGGAAGGAGATTGAAAGGACCTGGGAAGAGGTTCATTGGTCATGGTGCCTGGCTGGTGCTTTCCAAAGCCAAGGTTGCTGGTGTGTGCCTGACAGTTCCATAGGAAGTAGGGACTTCTGGGAGGGAGGAGGAGGTGGAAGGTGCCCTG - 3'

### Supplementary Table 2 | Transfection and selection details for each experiment

| | | | Step 1 - Deletion | | | | | | | Step 2 - Reinsertion | | | | | | |
|---|---|---|---|---|---|---|---|---|---|---|---|---|---|---|---|---|
| **Locus** | **Target(s)** | **Cell Line** | **sgRNAs** | **Cell Density (cells/well)** | **DNA added** | **Incubation time** | **SF added or exchanged** | **Time puro added** | **[Puro]** | **Hours before transfection** | **Cell Density (cells/well)** | **DNA added** | **Incubation time** | **SF added or exchanged** | **Time puro added** | **[Puro]** |
| *TSNARE1* | rs4129585 | RU04 | 1 and 3 | 40K | 500ng | 7 hours | Added | ~48 hours | 1ug/mL | 24 | 40K | 500ng | 7 hours | Added | ~40- 48 hours | 1 ug/mL |
| *TSNARE1* | rs4129585 | RU17 | 1 and 3 | 40K | 500ng | 6 hours | Exchanged | 24-48 hours | 0.5ug/mL - 1ug/mL | 24 | 40K | 500ng | 6 hours | Added | ~46 hours | 1.20ug/mL |
| *NRGN* | rs12293670 | RU02 | 1 and 2 | 40K | 500ng | 4 hours | Added | 40 hours | 1.2 ug/mL | 36 | 30K | 440ng | 14hrs | Exchanged | 40 hours | 1.5ug/mL |
| *ATP2A2* | rs4766428 | RU07 | 4 and 2 | 30K | 500ng | 4 hours | Added | 48 hours | 1.5ug/mL | 24-48 | 40-50k | 500ng | 5-7 hours | Exchanged | 24-48 hours | 1 ug/mL |
| | | RU02 | 4 and 2 | 50k | 500ng | 4 hours | Added | 24 hours | 1ug/uL | 120 | 30k | 585ng | 6-8 hours | Exchanged | 24 hours | 1ug/mL |
| | | RU05 | 4 and 2 | 30K | 500ng | 4 hours | Added | 24 hours | 1.5ug/mL | N/A | N/A | N/A | N/A | N/A | N/A | N/A |
| | | RU05 | 1 and 2 | 30K | 500ng | 4 hours | Added | 24 hours | 1.5ug/mL | N/A | N/A | N/A | N/A | N/A | N/A | N/A |
| *NRXN1* | Exon 2 (ENSE00002171876) | RU02 | 9 and 10 | 30K | 500ng | 4 hours | Exchanged | 24 hours | 1.3ug/mL | N/A | N/A | N/A | N/A | N/A | N/A | N/A |
| *DPYSL2* | Exon 1 (ENSE00002094501) | RU13, RU03 | 1 and 3 | 30K | 500ng | 4 hours | Added | 48 hours | 1ug/mL | N/A | N/A | N/A | N/A | N/A | N/A | N/A |
| *PKNOX2* | Exon 3 (ENSE00002153508) | RU05 | 1 and 2 | 55K | 500ng | 4 hours | Added | 24 hours | 1 ug/mL | N/A | N/A | N/A | N/A | N/A | N/A | N/A |

### Supplementary Table 3 | Summary of Off-Target Analyses

| | | | **Step 1 - Deletion** | | | | | **Step 2 - Reinsertion** | | | | |
|---|---|---|---|---|---|---|---|---|---|---|---|---|
| **Locus** | **Target(s)** | **sgRNAs** | **Sites Screened** | **Clones Screened** | **Positive Clones** | **Positive Sites** | **Edits/Possible Edits** | **Sites Screened** | **Clones Screened** | **Positive Clones** | **Positive Sites** | **Edits/Possible Edits** |
| *TSNARE1* | rs4129585 | 1 and 3 | 4 | 2 (RU04)<br>2 (RU17) | 0 | 0 | 0/16 | 8 | 10 (RU04)<br>10 (RU17) | 2 (RU17)<br>3 (RU04) | 2 | 5/160 |
| *NRGN* | rs12293670 | 1 and 2 | 14 | 1 | 0 | 0 | 0/14 | 0 | 0 | 0 | 0 | |
| *ATP2A2* | rs4766428 | 4 and 2 | 9 | 3 (RU07)<br>2 (RU02) | 0 | 0 | 0/45 | 17 | 11 (RU07)<br>6 (RU02) | 0 | 0 | 0/289 |
| | | 1 and 2 | 13 | 10 | 0 | 0 | 0/130 | N/A | N/A | N/A | N/A | |
| *NRXN1* | Exon 2 (ENSE00002171876) | 10 and 9 | 2 | 6 | 0 | 0 | 0/12 | N/A | N/A | N/A | N/A | |
| *DPYSL2* | Exon 1 (ENSE00002094501) | 1 and 3 | 1 | 6 | 0 | 0 | 0/6 | N/A | N/A | N/A | N/A | |
| *PKNOX2* | Exon 3 (ENSE00002153508) | 1 and 2 | 4 | 11 | 0 | 0 | 0/44 | N/A | N/A | N/A | N/A | |
| Deletion Off-Target Edits/Possible Edits | | | | | | | 0/267 | Reinsertion Off-Target Edits/Possible Edits | | | | 5/449 |
| **Total Off-Target Edits/Possible Edits** | | | | | | | | | | | | **5/716 (0.7%)** |

**Supplementary Table 4 | Off-Target sites and sanger sequencing primers**

| **Experiment: *TSNARE1* Deletion** | | |
|---|---|---|
| **Off target site** | **Forward primer** | **Reverse primer** |
| chr12: 104496044-104496066 | 5'-TGGAGTTCCTTGAAGTGCCCA-3' | 5'-TGTGGCCAGCAGTCACTTCA-3' |
| chr14: 99036175-99036197 | 5'-CCTCCCGCCTGTTTCTCATT-3' | 5'-TAACGCACTCAAGGCTGTGT-3' |
| chr7: 42054426 - 42054448 | 5'-TCCTCTCAGGGATAAGGCCC-3' | 5'-GCAGTCATGCAGAGATGCAG-3' |
| chr8: 129272886- 129272908 | 5'-CCACGGGGAGAACAGAACTC-3' | 5'-GCTCCGTAAACATAACGCCG-3' |

| **Experiment: *TSNARE1* Re-insertion** | | |
|---|---|---|
| **Off target site** | **Off target site** | **Off target site** |
| chr20:41716915-41716937 | chr20:41716915-41716937 | chr20:41716915-41716937 |
| chr6:158640569-158640591 | chr6:158640569-158640591 | chr6:158640569-158640591 |
| chr19:1423000-1423022 | chr19:1423000-1423022 | chr19:1423000-1423022 |
| chr5:52639468-52639490 | chr5:52639468-52639490 | chr5:52639468-52639490 |
| chr1:155240834-155240856 | chr1:155240834-155240856 | chr1:155240834-155240856 |
| chr12:58096131-58096153 | chr12:58096131-58096153 | chr12:58096131-58096153 |
| chr18:7231329-7231351 | chr18:7231329-7231351 | chr18:7231329-7231351 |
| chr10:102402661-102402683 | chr10:102402661-102402683 | chr10:102402661-102402683 |

| **Experiment: *ATP2A2* guides 1&2 Deletion** | | |
|---|---|---|
| **Off target site** | **Forward primer** | **Reverse primer** |

| | | |
|---|---|---|
| chr1:64608484-64608506 | 5'-CCCACACACACACTTTCACC-3' | 5'-TGGGTCCCTGTTACATCCAT-3' |
| chr5:70203940-70203962 | 5'-TTGGAAAACCTGGGTGCTAC-3' | 5'-CACCTGGCCCTTTAACTTGA-3' |
| chr5:69328521-69328543 | 5'-TTGGAAAACCTGGGTGCTAC-3' | 5'-CACCTGGCCCTTTAACTTGA-3' |
| chr15:50458632-50458654 | 5'-TCCAGCAGGAGGAAAATGTC-3' | 5'-TGCTTTGGCTCCCTGTAGAC-3' |
| chr7:104614660-104614682 | 5'-TCACGTGTCCCAAATGTGTT-3' | 5'-CCCAACTCAGTCAGCCCTAA-3' |
| chr10:16731210-16731232 | 5'-GTGTGCCTGTAGTCCCAGGT-3' | 5'-ACAAGGCTAGGGCATTTCAA-3' |
| chr13:26322031-26322053 | 5'-CAGGATGTCAGGATGTGTGC-3' | 5'-CTGTTTTGGAAACCCAGAGC-3' |
| chr12:39537192-39537214 | 5'-CACCACGTGGAACTGTTGAG-3' | 5'-TTGGTAAGGGTGGAGGTTTG-3' |
| chr4:170035824-170035846 | 5'-AGGTCCTTTTGGCCATGTAA-3' | 5'-AAGGGAGCCTGTAAACAGCA-3' |
| chr16:51531515-51531537 | 5'-TTGCATAGTGGCGAAGTCAG-3' | 5'-CTGGAGGTGAGTCCAACCAT-3' |
| chr5:90235329-90235351 | 5'-AGGCTCACAGCTGACTTTCC-3' | 5'-AATGGCATCAGGTTTGAAGG-3' |
| chr13:45417817-45417839 | 5'-CAGACAAGGCCACTCTGTGA-3' | 5'-GCAAAGACCACCAGGAACAT-3' |
| chr8:117277149-117277171 | 5'-CTGCCCAAGGTCATTTGTCT-3' | 5'-CCAGGGATTCCATCAACCTA-3' |

| **Experiment: *ATP2A2* guides 4&2 Deletion** | | |
|---|---|---|
| **Off target site** | **Forward primer** | **Reverse primer** |
| chr6:157351760-157351782 | 5'-CACCTGGTCACCCAAGAACT-3' | 5'-ACTTGCAGTTCCTTCCTCCA-3' |
| chr2:134135281-134135303 | 5'-GTGGTCTGGCCCAGCTAAT-3' | 5'-GCCAGGATGGTCTCAATCTC-3' |
| chr7:133626127-133626149 | 5'-GCAGGTGGATCACTTGAGGT-3' | 5'-GTAAAGGTCACGGCCAAAAA-3' |
| chr13:45417817-45417839 | 5'-GCAAAGCCAAAAGCAAGATT-3' | 5'-TTCCAGGGTCATTCAGATCC-3' |
| chr1:187899177-187899199 | 5'-TTGCCAAAGATGTTCAGAGC-3' | 5'-CCTTCCTTGCCAGTTGTGTT-3' |
| chr8:37542827-37542849 | 5'-AGTCTCGCTGTTGTCCACCT-3' | 5'-TGACCCTGGGTGTACATTGA-3' |
| chr8:117277149-117277171 | 5'-TCAACAGAGAAAATGCTTTGGA-3' | 5'-CCCTCAAGAAGGATGGAACA-3' |
| chr10:45021897-45021919 | 5'-AGCCCCCTAGGTTCTCAGAC-3' | 5'-GCTGGAGTGCAGTGGTGTAA-3' |

| | | |
|---|---|---|
| chr13:114811452-114811474 | 5’-GGAAAGACCCAAGCGTGTAA-3’ | 5’TTTTCTTCCACCACCACCTC-3’ |

| **Experiment: *ATP2A2* Re-insertion** | | |
|---|---|---|
| **Off target site** | **Forward primer** | **Reverse primer** |
| chr2: 82781444-82781466 | 5'-CCCTTCCAGCTGTCCACTTA-3' | 5'-GTTTTTAGCGCTGTCCGTTT-3' |
| chr12:16732215-16732237 | 5'-AATGGTTTGGCTGTTTTCCA-3' | 5'-CAGCCTCAGCCTTCCTACTG-3' |
| chr9:79893486-79893508 | 5'-TTGCTTTGCCAAGAACAAAGGAT-3' | 5'-GCTTCTGCTCAAAATGTGCT-3' |
| chr13:108916590-108916612 | 5'-TCTTGCCGTCACTGTTTCAG-3' | 5'-TGTAGGCTCTGAAGCCAGAAG-3' |
| chr10:16731210-16731232 | 5'-GTGTGCCTGTAGTCCCAGGT-3' | 5'-ACAAGGCTAGGGCATTTCAA-3' |
| chr4:136106449-136106471 | 5'-AAGTCTGGGTGCCAAAATGT-3' | 5'-GTCCCATGTTGCTAGGAAGG-3' |
| chr1:77027949-77027971 | 5'-CCTTGGGCTGATGAAAATGT-3' | 5'-GGAGATGGTGGACAGACGTT-3' |
| chr6:75881598-75881620 | 5'-AGAAAAACAGCAGCCCTCAA-3' | 5'-CACACGCACGTGTCTTTGAT-3' |
| chr13:45417817-45417839 | 5'-AGCAAAGTGGCACCATTACC-3' | 5'-CAAAGTTGGGTGGGTTGACT-3' |
| chr18:24305713-24305735 | 5'-CCGGGATGAATTTTACTTGG-3' | 5'-TCCTTGAGATGCAAACCTCAT-3' |
| chr11:82686439-82686461 | 5'-AAGCCTAAACCGGCCTTAAA-3' | 5'-AAATGGAGGCATGAATCTCG-3' |
| chr17:12980226-12980248 | 5'-CTGGCATGCCCTAAGAACTC-3' | 5'-ACAGGCGAGCAGCATAAGAT-3' |
| chr8:74630198-74630220 | 5'-TGGCTCCCACACTGTTCTTT-3' | 5'-ACCAGGAGGCTCTGATATGC-3' |
| chr16:51531515-51531537 | 5'-TTGCATAGTGGCGAAGTCAG-3' | 5'-GAAATAGCCCAGGCAAGACA-3' |
| chr8:117277149-117277171 | 5'-CTGCCCAAGGTCATTTGTCT-3' | 5'-CCAGGGATTCCATCAACCTA-3' |
| chr4:170035824-170035846 | 5'-AGGTCCTTTTGGCCATGTAA-3' | 5'-AAGGGAGCCTGTAAACAGCA-3' |
| chr12:15851243-15851265 | 5'-CTTGGCCAAATGGAGATGTT-3' | 5'-GGCAGGGAAAGTCATCATGT-3' |

| **Experiment: *NRXN1* Deletion** | | |
|---|---|---|
| **Off target site** | **Forward primer** | **Reverse primer** |

| | | |
|---|---|---|
| chr13:112328963-112328985 | 5’-AGTTCAGCATAGCCCAACCA-3’ | 5’-CCTGACCCAAATGCTACAGTC-3’ |
| chr14:93417365-93417387 | 5’-TCTCTAGGCTGACCCCTTCC-3’ | 5’-GTTTGCCCGAAGTCCAGAGA-3’ |

| **Experiment: *DPYSL2B* Deletion** | | |
|---|---|---|
| **Off target site** | **Forward primer** | **Reverse primer** |
| chr14:67889176-67889198 | 5’-CAGGCAACTAGGGACCACTC-3’ | 5’-GCCCAACTTCTTTGTGGAAA-3’ |

| **Experiment: *NRGN* Deletion** | | |
|---|---|---|
| **Off target site** | **Forward primer** | **Reverse primer** |
| chr9:73488609-73488631 | 5’-CTTTACGCTCCAGCAAACCA-3’ | 5’-ACTTCAGCCCACAAAGAGATTA-3’ |
| chr2:99389481-99389503 | 5’-GGACAACCTACTCCTCCAGC-3’ | 5’-GAATCACCACACACCCTCCA-3’ |
| chr19:51783617-51783639 | 5’-ACAACATGACCCAGATCACAAGA-3’ | 5’-CAGACCCACCTAGCATACTCC-3’ |
| chr9:22117784-22117806 | 5’-TCATTTGGAAGCTCGTGGGA-3’ | 5’-GATCCAAGCGCTACCCATCC-3’ |
| chr18:61557728-61557750 | 5’-CGCAAGAAAGAAGCGACCAAT-3’ | 5’-CCAGGCAAGATTGCAGGGATA-3’ |
| chr21:45614885-45614907 | 5’-GACCCCAACACAAGGAGGAC-3’ | 5’-GCAAATCTGCTTGAGGCTCG-3’ |
| chr6:143091823-143091845 | 5’-AGCCACATGGGGGTGGAAAA-3’ | 5’-CCCCCACTCCTGCCTGTATT-3’ |
| chr11:92710875-92710897 | 5’-GTACTTCTCCAAGCCCGCAT-3’ | 5’-GCTCAGCACCCTCAGAAGAG-3’ |
| chr5:43675682-43675704 | 5’-AAATGCCCATCCCCAGAACA-3’ | 5’-GGACAGGCTACCTCCCCTTA-3’ |
| chr16:68420724-68420746 | 5’-GTGTGCAATTGATTTGATAGGGAA-3’ | 5’-ATCTTGACAACGGGCCCAAAA-3’ |
| chr22:49146042-49146064 | 5’-AGCCCATGAGGAAAGGCTTG-3’ | 5’-CTGACCTGCTCCCCCTTAGA-3’ |
| chr11:100901165-100901187 | 5’-CTCGGACTTCACCCGTTCTC-3’ | 5’-ACAGAGGAATAAAGCGGCCT-3’ |
| chr14:77795643-77795665 | 5’-GATTCCCTGCATCTCTTGCC-3’ | 5’-TTGCTATGTGCTGGAGGCTT-3’ |
| chr5:174771326-174771348 | 5’-TTGGTTGAAGGAAGGGGCTC-3’ | 5’-GCTCTCACTCTGATGGCTGG-3’ |

| **Experiment: *PKNOX1* Deletion** |
|---|

| Off target site | Forward primer | Reverse primer |
|---|---|---|
| chr11: 58525010-58525032 | 5’-GCTCAGGGAGTCACTTGCTT-3’ | 5’-GAATCACCCTGGAAATGGAA-3’ |
| chr9:112952518-112952540 | 5’CAGCCCCATGCGTATTTTAT-3’ | 5’GGAGTTTCTTGCCCAAATCA-3’ |
| chr7:73818508-73818530 | 5’-CAGAAGGTGGGGACAAGAAG-3’ | 5’-CGGGACTGTCACTTTGGAGA-3’ |
| chr16: 47533784-47533806 | 5’-GAGCAGTCATAGGTCGCACA-3’ | 5’-AGGGTGGATTCCCACTAAGG-3’ |

**Supplementary Table 5 | Cell line information**

| LAB_ID | Rutgers UID | Passage number upon receipt | Sex | Population | Designation |
|---|---|---|---|---|---|
| RU01 | MH0180967 | 10 | male | European | control |
| RU02 | MH0180966 | 12 | female | European | control |
| RU03 | MH0180968 | 11 | female | European | control |
| RU04 | MH0180965 | 12 | male | European | control |
| RU05 | MH0180971 | 11 | female | European | control |
| RU06 | MH0185943 | 19 | female | European | control |
| RU07 | MH0185941 | 7 | male | European | control |
| RU08 | MH0185940 | 7 | male | European | control |
| RU09 | MH0185933 | 12 | male | African | control |
| RU10 | MH0185938 | 9 | male | European | control |
| RU11 | MH0185917 | 9 | female | European | control |
| RU12 | MH0185910 | 11 | Female | European | control |
| RU13 | MH0185922 | 7 | male | African | control |
| RU14 | MH0185905 | 7 | female | European | control |
| RU15 | MH0185860 | 8 | male | European | control |

| | | | | | |
|---|---|---|---|---|---|
| RU16 | MH0185857 | 11 | Male | European | control |
| RU17 | MH0185863 | 7 | female | European | control |

**Supplementary Table 6 | Filtering pipeline for candidate off-target sites**

| **Experiment** | **Total Off-targets (up to 4 mismatches)** | **# with filtered mismatches (mismatches included)** | **Sites screened (in exons or overlapping open chromatin (distance from peak))** |
|---|---|---|---|
| rs4766428 *(ATP2A2)* 1+2 Deletion | 662 | 662 (1-4) | 13(0bp) |
| rs4766428 *(ATP2A2)* 4+2 Deletion | 901 | 14 (1-2) | 9(500bp) |
| rs4766428 *(ATP2A2)* 4+2 Reinsertion | 855 | 144 (1-3) | 17(0bp) |
| *DPYSL2B* Exon 1 Deletion | 45 | 1(1-3) | 1(500bp) |
| rs4129585 (*TSNARE1*) Deletion | 109 | 14 (1-3) | 4 (100bp) |
| rs4129585 (*TSNARE1*) Reinsertion | 88 | 7 (1-3) | 8 (500) |
| *NRXN1* Exon 2 Deletion | 112 | 3 (1-3) | 2(200bp) |
| rs12293670 (*NRGN*) Deletion | 404 | 44(1-3) | 14(0bp) |
| rs12293670 (*NRGN*) Reinsertion | 134 | 12(1-3) | 6(0bp) |
| *PKNOX1* Exon 3 Deletion | 347 | 26(1-3) | 4(300bp) |
